# Tools for accurate post hoc determination of marker location within whole-brain microscopy images

**DOI:** 10.1101/2021.05.21.445133

**Authors:** Adam L. Tyson, Mateo Vélez-Fort, Charly V. Rousseau, Lee Cossell, Chryssanthi Tsitoura, Horst A. Obenhaus, Federico Claudi, Stephen C. Lenzi, Tiago Branco, Troy W. Margrie

## Abstract

To interpret in vivo experiments designed to understand brain function, high-resolution whole-brain microscopy provides a means for post hoc determination of the location of implanted devices and recorded cells in three dimensional brain space that is a critical step for data interrogation. Here we have developed Python-based tools (brainreg and brainreg-segment) to accurately map, in a common coordinate space, the position of dye-labelled probe tracks and two-photon imaged cell populations expressing fluorescent protein. The precise location of probes and cells were validated using physiological recordings and human raters that indicate accuracy levels to less than 70µm. These flexible, open-source methodologies are expected to further evolve with need and to deliver the anatomical precision that is necessary for understanding the functional architecture of the brain.

## Introduction

The brain is a complex organ with many different macroscopic structures, defined by function, anatomy, connectivity and gene expression. To understand its function, modern systems neuroscience relies on the use of implanted devices for stimulating and recording neurons (Adelsberger et al., 2005; Arenkiel et al., 2007; Jun et al., 2017) and viral injections are often used to induce expression of proteins required, for e.g. imaging (Grienberger and Konnerth, 2012) or optical stimulation (Mei and Zhang, 2012) of genetically-defined neuronal populations. To correctly interpret the resultant data, it is critical to map the location of the implanted devices or injections. This allows for the brain region of interest to be defined in single animals, but more importantly, it allows the data from multiple animals to be pooled and compared in a common anatomical coordinate system. High-density electrophysiological probes such as Neuronexus (Berényi et al., 2014) and Neuropixels (Jun et al., 2017, Steinmetz et al., 2021) present additional challenges (Steinmetz et al., 2018). These probes consist of many hundreds of sites that can record activity in cells across multiple regions of the brain. Unlike other implanted devices, such as optical fibres, it is critical to locate the tip of the probe, as well as regions along the shank to ensure that recording sites can be assigned to specific regions and populations of cells.

Neuronal populations targeted by viral injection are often visualised by co-expression of a fluorescent protein. Implanted devices can either be localised by the “lesion track” left in the brain following their removal (e.g. optical fibers) or by coating the device with a fluorescent dye. Traditionally, mapping of these bulk signals in the rodent brain was carried out by tissue sectioning, microscopy and manual (Cooper et al., 1953; Renshaw et al., 1940; Starr et al., 1973) or semi-automated (Shamash et al., 2018, Peters, 2021) analysis. Analysis of two-dimensional tissue sections has a number of drawbacks, particularly when mapping large three-dimensional (3D) structures that appear in subsequent tissue sections. Processing, imaging, and analysing a series of images can be laborious, and computational reconstruction of these images into a 3D volume can introduce artefacts. Using block-face imaging methods (Gong et al., 2013; Ragan et al., 2012; Seiriki et al., 2017) or a combination of tissue clearing and light-sheet fluorescence microscopy (LSFM; Chung et al., 2013; Dodt et al., 2007; Gong et al., 2013; Ragan et al., 2012; Renier et al., 2014; Seiriki et al., 2017; Susaki et al., 2014), fully automated, cellular resolution imaging of whole, intact rodent brains is now possible (Osten & Margrie, 2013). Commercially available microscopes, and promising open-source initiatives (Campbell, 2020a; Economo et al., 2016; Tomer et al., 2014; Voigt et al., 2019) are leading to the rapid adoption of these methods.

The development of detailed open-source 3D brain atlases, such as the Allen Mouse Brain Common Coordinate Framework version 3 (CCFv3, Wang et al., 2020), means that brain images can be warped to an atlas image. This allows both the assignment of a detected feature (e.g. implanted device or injection volume) to a brain region, and also for the image to be warped to a common coordinate system, allowing for analyses of groups of animals in a standardised manner. There exist many methods for registering 3D whole-brain microscopy data to an atlas (Furth et al., 2018; Goubran et al., 2019; Jin et al., 2019; Ni et al., 2020; Niedworok et al., 2016; Renier et al., 2016; Young et al., 2020), but these are relatively inflexible, only supporting one atlas at a single resolution (Tyson and Margrie, 2021). This is problematic, as it does not allow researchers to use more recently developed atlases (Chon et al., 2019; Hoops et al., 2021; Kenney et al., 2021; Perens et al., 2020; Young et al., 2021), nor adapt software developed for one model organism, to another. Tools also exist for the detection and analysis of structures in whole-brain images such as neuronal somata (Furth et al., 2018; Goubran et al., 2019; Iqbal et al., 2019; Mano et al., 2020; Renier et al., 2016; Song et al., 2020; Tyson et al., 2020; Young et al., 2020), axons (Friedmann et al., 2020; Goubran et al., 2019) and vasculature (Kirst et al., 2020; Todorov et al., 2020). While mapping implanted devices within the brain has been performed using non-invasive magnetic resonance imaging (MRI) or computed tomography (CT) (Borg et al., 2015, Rangarajan et al., 2016, Király et al., 2020; Kollo et al., 2020), there has been only one study using 3D whole brain microscopy (Liu et al., 2020). Although conceptually simple, there exist no open-source, user-friendly tools for the detection and analysis of large structures in 3D whole-brain microscopy images.

Here, we address these problems with two integrated open-source software tools. The first is “brainreg”, a python port of the validated aMAP (Niedworok et al., 2016) registration and segmentation software. Brainreg integrates with the BrainGlobe Atlas API (Claudi et al., 2020) to allow registration of whole-brain microscopy images at multiple resolutions with different atlases. The use of the BrainGlobe Atlas API allowed brainreg to be developed in an atlas-agnostic manner, facilitating use with different atlases, including those yet to be developed. The second is “brainreg-segment”, a tool for manual segmentation of large structures, following registration, developed as a plugin for napari, a powerful new multidimensional image viewer, built to leverage the modern scientific Python ecosystem (Sofroniew et al., 2021). As brainreg and brainreg-segment are built upon the BrainGlobe Atlas API, they are interoperable with other tools in the suite, such as cell detection in cellfinder (Tyson et al., 2021) and, importantly, brainrender (Claudi et al., 2021), for visualisation of segmented structures in atlas space. In addition, we demonstrate the applicability of these two integrated open-source software tools by segmenting the geometric location of high-density silicon probe electrode sites and brain regions that contain fluorescent cells following viral injections.

## Results

### Image registration using brainreg

While there exist many excellent 3D whole-brain registration packages, they are at present limited in their compatibility with a single atlas. This places constraints on the animal model species that can be used, the analyses that can be performed, and also the computing hardware that must be used, as many atlases have very large file sizes. To overcome these issues, we developed brainreg, which is an updated version of aMAP that was validated by comparing its performance to that of experts performing manual segmentation. Compared to aMAP, brainreg is developed in Python, which is emerging as the programming language of choice in neuroscience (Muller et al., 2015), and it does not require any data preprocessing (e.g. downsampling). This simplifies both use, and potential further development within the community. Lastly, by using the BrainGlobe Atlas API, brainreg is not specific to any one atlas, and so the registration can be performed for data from different species, using atlases with different annotations, or at different resolutions to decrease processing time.

Brainreg loads the reference image volume from the chosen atlas (Figure 1A), and the raw data (Figure 1B), which is typically an autofluorescence channel. The image data is resampled and reoriented to match the atlas file, and then both images are filtered to remove-high frequency noise. The two images are firstly aligned using affine registration, and then by a non-linear free-form registration step. The resulting transformations are used to warp the raw data (and any other fluorescence channels within the same dataset, such as those containing cell- or region-specific labelling) into the coordinate space of the atlas (Figure 1C). In addition, atlas files (e.g. the CCFv3 annotations, Figure 1A) are also warped to the coordinate space of the raw data to allow analysis in either (raw, or atlas) coordinate spaces.

**Figure 1:**
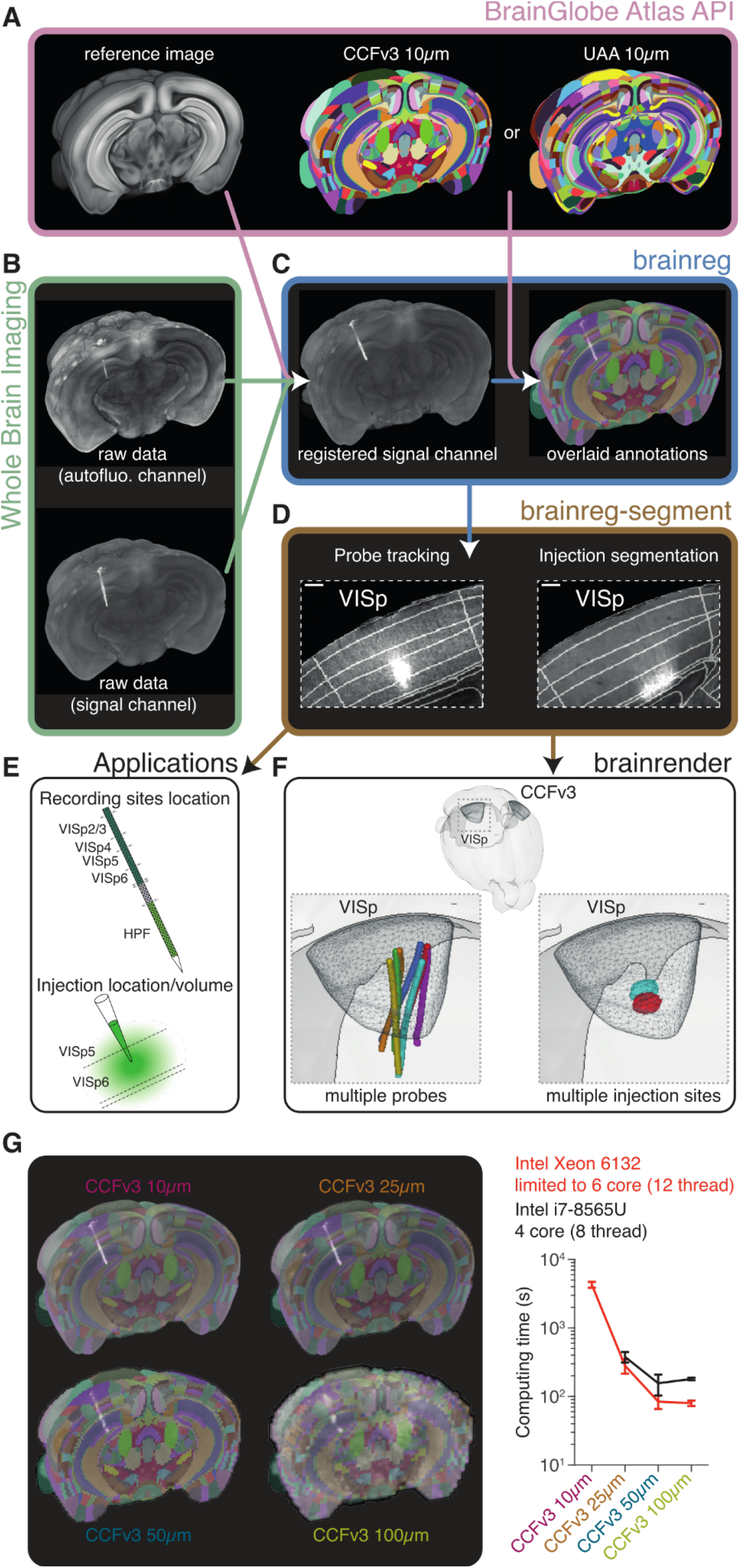
Overview of the brainreg and brainreg-segment workflow. **A**. Atlas data including the reference anatomical image and the brain region annotations are sourced from the BrainGlobe Atlas API and used for image registration. Multiple atlases, such as the Allen Mouse Brain Common Coordinate Framework version 3 (CCFv3) and the Unified Anatomical Atlas (UAA) are supported. **B**. Multichannel (e.g. labelled signal channel and a secondary autofluorescence channel) whole-brain microscopy data can be processed directly by brainreg without preprocessing (other than stitching). **C**.Brainreg registers the sample data to the atlas reference image (and vice versa), allowing brain region annotations to be overlaid upon the raw data. **D**. Brainreg-segment allows manual segmentation of labelled structures such as linear electrophysiological probes and fluorescent cell populations. **E**.Once segmented, the anatomical distribution of labelled structures can be analysed, for e.g. by assigning probe recording sites or cellular distribution to a cortical layer. **F**. Segmented structures from multiple animals can be directly exported to brainrender for visualisation in a common anatomical space (along with other data, such as atlas brain regions). Probes and injection sites visualized in Allen CCFv3 space, along with the target brain region, primary visual cortex (VISp, wireframe). **G**. Registration of a single brain to the CCFv3 at 10, 25, 50 and 100 μm isotropic resolutions.m isotropic resolutions. Registration on a desktop workstation (Intel Xeon 6132, red) at 10 μm took 4283 ± 425 seconds, but all other resolutions are much quicker (<281 ± 66 seconds). Registration at 25, 50 and 100 μm on a laptop (Intel i7-8565U, black) was slower, but still relatively fast (<378 ± 67 seconds).

To segment bulk structures such as labelled electrophysiological probes or the spread of viruses encoding fluorescent proteins, an additional tool called brainreg-segment was developed (Figure 1D). Briefly, the user is shown the results of the registration step (e.g. downsampled data and atlas annotations), and they can segment (manually annotate) either 1D tracks, 2D areas, or 3D volumes. These segmented structures are analysed to determine their position and spatial extent in the brain, such as the brain region associated with each recording site on a probe, or the volume and spatial location of an injection (Figure 1E). The segmented structures can be exported as meshes to brainrender to allow for easy visualisation in atlas space, along with other structures, such as atlas brain regions (Figure 1F). For more details on the user interface, see the documentation (docs.brainglobe.info/brainreg-segment).

Brainreg retrieves atlas data from the BrainGlobe Atlas API (Figure 1A), which means that in addition to using different brain region annotations (see below), it can also be used to register data at different resolutions. Some atlases, such as the CCFv3, are available at different resolutions (10, 25, 50 and 100 μm isotropic, Figure 1G). Registration using high-resolution atlases can take a long time, and require specific computing resources (e.g. sufficient memory). To test the computational hardware required by brainreg, we tested the registration speed on two datasets using the CCFv3 five times at each resolution. We firstly tested the pipeline using a high-specification image analysis workstation (Figure 1G). Registration using the 10 μm atlas took over an hour (4283 ± 425 seconds), but all other resolutions took less time (<281 ± 66 seconds). However, researchers may not have access to powerful analysis workstations, so we repeated these tests on a standard laptop. The 10 μm atlas could not be used, as the memory required (∼30GB) was not available, but registration using the other atlases was fast (<378 ± 67 seconds).

### Localization of silicon probes

To demonstrate the advantages of the analysis pipeline and its applications, we assessed the performance of 3 neuroscientists in localising silicon probe tracks in 7 imaged mouse brains. Briefly, a high-density silicon probe (Neuropixels) was coated with DiI and the tip inserted in the primary visual cortex 1,750 to 2,000 mm deep. After recordings, the animal was perfused, the brain harvested and imaged using serial two-photon tomography (Ragan et al., 2012; Figure 2A). A total of 7 brains (each with one silicon probe track) were imaged and then processed through the brainreg pipeline (Figure 1 A-C) using the CCFv3 10 µm resolution atlas (Figure 1G).

**Figure 2.**
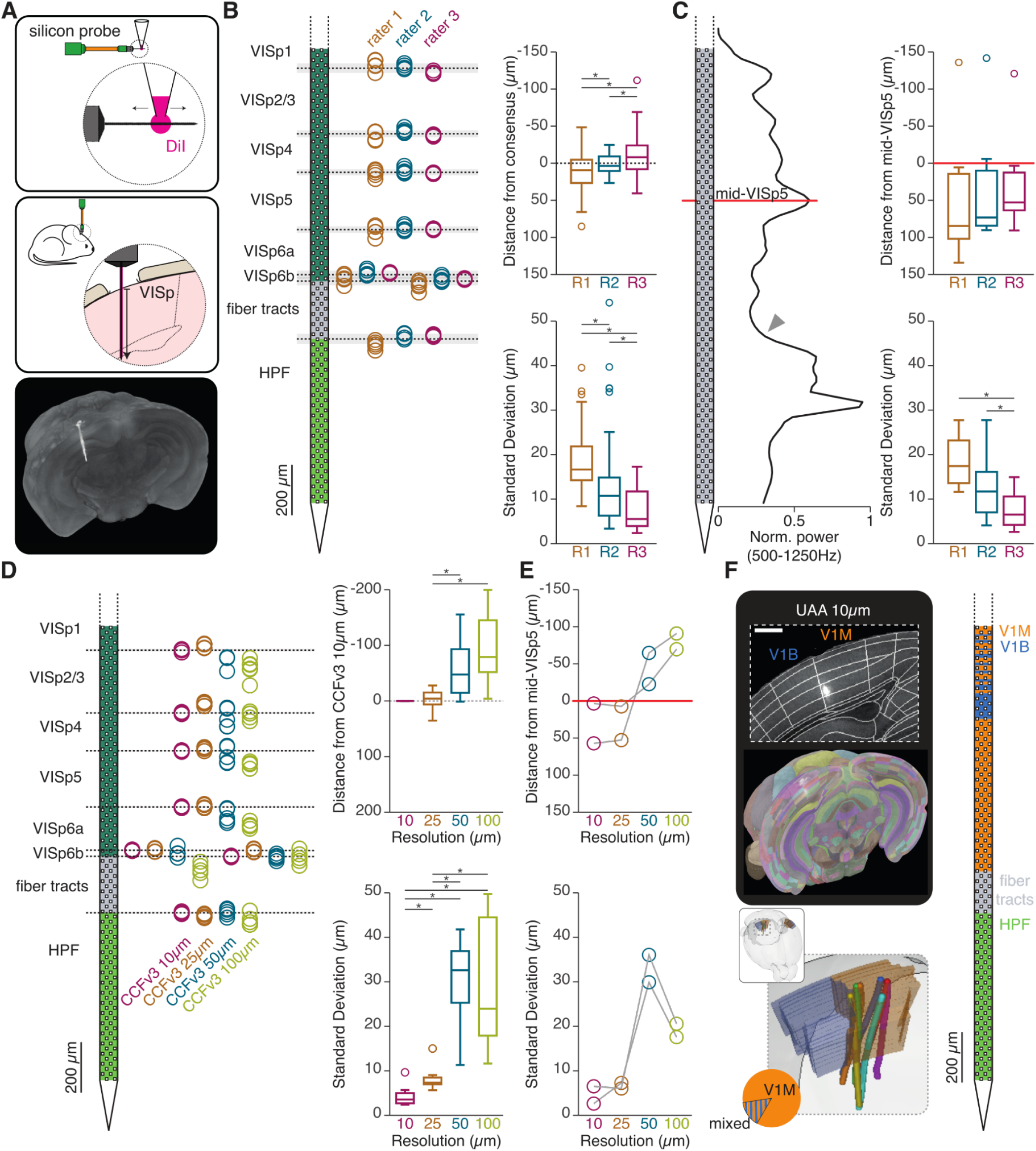
Accurate and reliable silicon probe track tracing with brainreg-segment. **A**. Schematic of the experiment timeline. Top: a Neuropixels probe is coated with DiI by coming into contact with a suspended drop of the dye. Middle: the Neuropixels probe is inserted in the mouse brain above the primary visual cortical region (VISp) at a depth of 1750 or 2000 μm. Bottom: Projection of the first 212 brain images (red channel) acquired with serial two-photon tomography, showing the DiI signal left by the Neuropixels probe. **B**. Left: Schematic of the probe and the position of each cortical layer crossing extracted from each raters’ track tracing (circles, n = 5 track tracings per rater). Plot of the average layer crossing depth for all raters (dashed black line) and standard deviation (shaded grey line). Top right: Box plot showing the distance from consensus for all cortical layers and for each rater. Bottom right: Box plot showing the standard deviation of each rater. **C**. Left: Schematic of the probe and plot of the normalized power against depth. The red line shows the depth of the cortical LFP power peak, indicating the position of the middle of layer 5 (mid-VISp5). The grey arrow indicates the increase in LFP power when entering the hippocampal formation. Top right: Box plot showing the distance from landmark (mid-VISp5) for each rater. Bottom right: Box plot showing the standard deviation of each rater. **D**. Left: Schematic of the probe and the position of each cortical layer crossing extracted from each CCFv3 atlas resolution (circles, n = 5 track tracings per atlas). Plot of the average layer crossing depth for the CCFv3 10 μm atlas (dashed black line) and standard deviation (very thin, shaded grey line). Top right: Box plot showing the distance from the CCFv3 10 μm atlas layer crossing (dashed grey line). Bottom right: Box plot showing the standard deviation for each atlas. **E**. Top: Plot showing the distance from landmark (mid-VISp5) for each brain and each atlas. Bottom: Plot showing the standard deviation for each brain and each atlas. **F**. Top left: coronal image of the primary visual cortex (inset) and projection of the first 223 brain images (red channel) acquired with serial two-photon tomography, overlaid with the UAA atlas. Scale bar = 500 μm. Right: schematic of the silicon probe and depth crossings from monocular primary visual cortex (V1M, orange) and binocular primary visual cortex (V1B, blue). Bottom left: brainrender rendering of the position of all the probe tracks (lines) relative to V1M (orange) and V1B (blue). Chart plot shows the probes that were exclusively located in V1M and those that were located in both V1M and V1B (mixed).

The 3 raters were then asked to localise the silicon probe track (DiI signal) and use the brainreg-segment “Track tracing” feature (Figure 1D) to record its position in the brain (brainreg-segment fitting properties: fit degree = 1; spline smoothing = 0.1; spline points = 1750 or 2000 [1 per µm]). In order to assess the performance of individual raters, each rater traced each probe track 5 times. Finally, for each tracing, cortical layer crossings were recorded thanks to the automated analysis feature of brainreg-segment and plotted according to depth (Figure 2B).

To assess the general variability of track tracing, a “consensus” layer crossing depth was computed by averaging recorded cortical depths across all cortical layers and across all raters (7 brains, 7 cortical layer crossings; n = 49 per rater). Then, each rater’s performance was compared to this general “consensus” by computing their average distance to the consensus (positive distances are closer to the pia; Figure 2B). The average distance from consensus was significantly different between raters, but was always below 10 µm (rater 1: median = 9.4 µm, range −48.4 to 85.2 µm; rater 2: median = 3.5 µm, range −24.7 to 26.6 µm; rater 3: median = −8.1 µm, range −111.8 to 40.6 µm; p<0.05 wilcoxon signed rank test). Excluding outliers, the maximum distance to consensus didn’t exceed +65.7 µm (rater 1) and −69.1 µm (rater 3). The outliers (2 out of 147 data points) represented a distance to consensus of +103.7 µm (rater 1) and −111.8 µm (rater 3; Figure 2B).

In order to assess the intra-rater probe tracing variability, we additionally computed the standard deviation of each rater’s layer crossing depths across brains (5 traced tracks in 7 brains with 7 cortical layer crossings; n = 49 SDs per rater). We found that reliability differed significantly between raters, but was always below an average of 17 µm SD (rater 1: median = 16.7 µm, range 8.4 to 39.5 µm; rater 2: median = 10.8 µm, range 3.4 to 54.2 µm; rater 3: median = µm, range 2.4 to 17.3 µm; p<0.05 wilcoxon signed rank test). Excluding outliers, the standard deviation never exceeded 31.9 µm (rater 1) and including outliers (7 out of 147 data points), it didn’t exceed 54.2 µm (rater 2; Figure 2B).

Overall, these results show that human track tracing using the brainreg/brainreg-segment pipeline was accurate. The average distance to the raters’ consensus varied significantly between raters, but without exceeding 10 µm on average and was typically below 70 µm. Human raters using the brainreg/brainreg-segment pipeline were also reliable, with an average intra-rater standard deviation of less than 17 µm and restricted to a typical maximum of 30 µm.

So far, the performance of track tracing has been compared to the raters’ general “consensus”. In extracellular recordings, however, neuronal activity patterns can, sometimes, allow experimenters to have one or several “ground-truth” depth data points. For instance, in the primary visual cortex, the depth distribution of local field potential power provides information about cortical layer localisation: a large-amplitude peak of the depth profile of power (500 to 1250 Hz) corresponds to mid-layer 5 (mid-VISp5, Figure 2C, Senzai et al., 2019). We used this prominent landmark to assess the raters’ accuracy and reliability against this “ground-truth” depth data point.

To assess the variability of track tracing, the mid-VISp5 location was computed from each rater’s traced tracks in each brain (7 brains, 1 mid-VISp5 crossing; n = 7 per rater). Then, each rater’s performance was checked by comparing their mid-VISp5 to the “ground-truth” mid-VISp5 computed from the LFP power profile (Figure 2C). The average distance from “ground-truth” (all raters: median = 57.4 µm, range = −141.7 to 134.1 µm) was not different between raters and was always below 84.5 µm (rater 1: median = 84.5 µm, range −135.9 to 134.1 µm; rater 2: median = 73.1 µm, range −141.7 to 90.3 µm; rater 3: median = 52.8 µm, range −120.8 to 90.6 µm; p>0.05 wilcoxon signed rank test). Excluding outliers, the largest distance to “ground truth” didn’t exceed +134.1 µm.

Interestingly, the distance from the landmark was positive in 94.4% of the cases when excluding outliers (17 out of 18 data points excluding outliers; range = 3.4 to 134.1 µm; only negative data point = −5.8 µm). These results suggest that there is a systematic underestimation of the probe track length, resulting in the “ground-truth” mid-VISp5 position being systematically shallower than the raters’ traced tracks (underestimation median = 69.4µm, range = −5.4 to 134.1 µm, n = 18, excluding outliers; Figure 2C; see discussion). When including outliers, the distance to the electrophysiological landmark didn’t exceed −141.7 µm (rater 2). However, the outliers in this data set (3 out of 21 data points) all originate from one single brain and each rater has it as an outlier (Figure 2C). Taking a closer view to this particular brain showed that the quality of the DiI labelling was suboptimal (due to brain surface deformation), indicating that the quality of the DiI tracing and brain tissue have a strong effect on track tracing accuracy and that errors introduced by suboptimal labelling are systematically reported by all raters.

In order to assess the intra-rater variability in localising mid-VISp5, we computed the standard deviation of each rater’s mid-VISp5 position across brains (5 traced tracks in 7 brains with 1 mid-VISp5 crossing; n = 7 SDs per rater). We found that reliability differed significantly between raters (with the exception of raters 1 and 2) and was always below an average of 17.4 µm SD (rater 1: median = 17.4 µm, range 11.6 to 27.7 µm; rater 2: median = 11.7 µm, range 4.1 to 27.8 µm; rater 3: median = 6.5 µm, range 2.6 to 14.9 µm; p<0.05 wilcoxon signed rank test). The raters’ standard deviations never exceeded 27.8 µm (rater 2; Figure 2C).

These results show that, when compared to an electrophysiological “ground-truth” landmark, the accuracy of all raters was on average 69.4 µm. The distance to the “ground truth” typically didn’t exceed 135 µm. The raters’ reliability was typically below a standard deviation of 17.4 µm.

After assessing raters performance in tracing probe tracks using the brainreg/brainreg-segment pipeline, the effect of atlas resolution was studied. Rater 3 was asked to trace probe tracks in 2 brains registered at 3 additional resolutions: 25, 50 and 100 µm (Figure 1G). Each track was traced 5 times, to assess the rater’s reliability (Figure 2D). The average distance of layer crossings was then compared to the layer crossing depths acquired by the same rater in the 10 µm resolution brain (2 brains, 7 cortical layer crossings; n = 14 per atlas). The distances recorded in the 25 µm resolution brains spread closely to the average layer crossings recorded in the 10 µm brains (median = −4.3 µm, range = −27.8 to 35.4 µm; Figure 2D). This distribution was in fact similar to that recorded in the 10 µm resolution brain (see rater 3, Figure 2B). However, the distances recorded in the 50 µm and 100 µm resolution brains were significantly greater (50 µm atlas: median = −47.5 µm, range = −155.4 to +1.2 µm; 100 µm atlas: median = −79.0 µm, range = −199.6 to −4.0 µm, respectively; p<0.05, wilcoxon rank sum test). In fact, in both brains recorded at a resolution of 50 or 100 µm resolution, the localisation of layer 1 (VISp1) and layer 6b (VISp6b), the thinnest of all cortical layers, were missed in several tracings. Interestingly, the accuracy deteriorated in one direction, indicating that lowering the brain resolution resulted in overestimating the probe length (see discussion).

In addition, the standard deviation of rater 3 (5 traces per brain, 2 brains, 7 cortical layer crossings; n = 14 SDs per atlas) were significantly different, but of the same order of magnitude for brains recorded at 10 or 25 µm (10 µm atlas: median = 3.5 µm, range = 2.4 to 9.6 µm; 25 µm atlas: median = 7.3 µm, range = 5.6 to 15.0 µm; p<0.05, wilcoxon signed rank test). However, the SDs significantly deteriorated in brains recorded at 50 and 100 µm (50 µm atlas: median = 32.6 µm, range = 11.3 to 41.8 µm; 100 µm atlas: median = 23.9 µm, range = 11.6 to 49.7 µm; p<0.05, wilcoxon rank sum test; Figure 2D), indicating that the reliability of track tracing deteriorated with lower resolutions. Similar results were found when comparing the distance to the “ground-truth” landmark: the tracing accuracy and reliability was similar between brains recorded at 10 and 25 µm, while they deteriorated when tracing in brains at resolutions of 50 and 100 µm (Figure 2E).

Overall, these results show that using the brain resolution of 25 µm has little effect in the localisation of probe tracks, while at the same time takes full advantage of processing speed (Figure 1G).

Finally, the traced probes by rater 3 were tested in the Unified Anatomical Atlas (UAA, Chon et al., 2019), which includes the border between the monocular and binocular primary visual cortex (V1M and V1B, respectively; Figure 1A). Out of the 7 probe tracks, only one was found to cross the V1M/V1B border (“mixed” V1M + V1B, Figure 1F), allowing the experimenter to identify which recordings originated exclusively from the monocular part of the primary visual cortex.

### Localisation of injections and functional characterisation of neurons

One useful application of the brainreg/brainreg-segment pipeline is the possibility of segmenting brain regions according to pathological irregularities, lesions or injections. We demonstrated this feature by asking 3 neuroscientists to manually segment a brain region where neurons expressed the green fluorescent protein (GFP).

Briefly, a glass pipette containing a solution of adenovirus carrying a “flexed” construct expressing GFP was inserted in the primary visual cortex of Ntsr1-cre transgenic mice. A small volume of the solution was injected at a depth of 900 µm (Figure 3A). Two to three weeks later, the animal was perfused, the brain harvested and imaged using serial two-photon tomography (Figure 3A). Two brains were imaged and then processed through the brainreg pipeline (Figure 1 A-C) using the CCFv3 25 µm resolution atlas (Figure 1G). Then, 3 raters were asked to segment the brain region containing GFP+ cells using the brainreg-segment “Region segmentation” feature (Figure 1D). In order to assess the performance of individual raters, each rater segmented each region 3 times.

**Figure 3.**
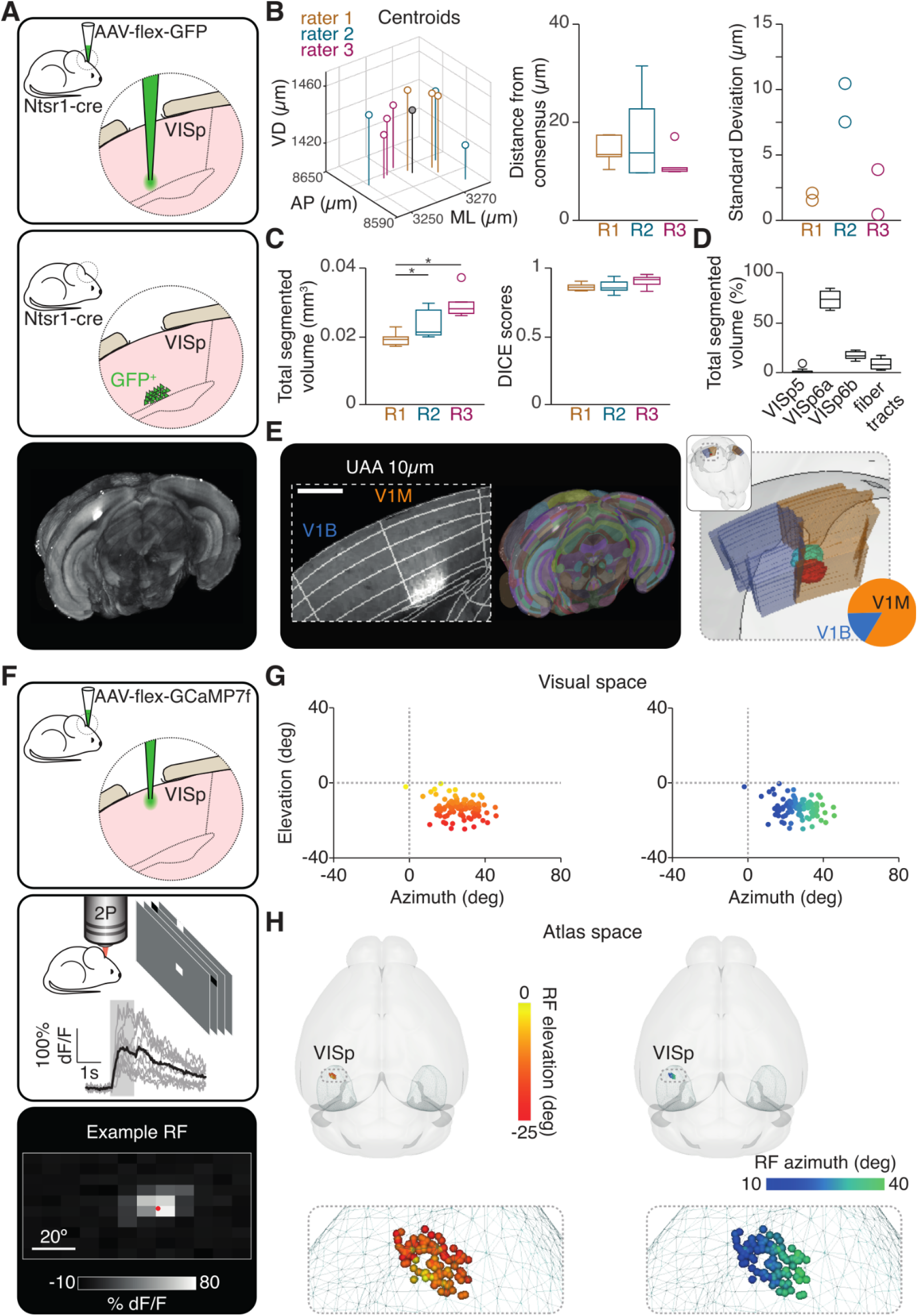
Accurate and reliable injection segmentation with brainreg-segment. **A**. Schematic of the experiment timeline. Top: Injection of an AAV carrying the fluorescent protein GFP in the primary visual cortex (VISp) at a depth 900 μm. Middle: Expression of GFP by a subset of cortical neurons. Bottom: Projection of the first 850 brain images (green channel) acquired with serial two-photon tomography, showing the GFP signal. **B**. Left: Plot showing the 3D position of the segmented region centroids by each rater (open circles, n = 3 centroids per rater). The average centroid for all raters is also shown (grey circle). Middle: Box plot showing the distance from consensus for all centroids and for each rater. Right: Plot showing the standard deviation of each brain and for each rater. **C**. Left: Box plot showing the total segmented volume for each rater. Right: Box plot showing the DICE scores for each rater. **D**. Box plot showing the percentage of segmented volume in each CCFv3 brain region. **E**. Left: coronal image of the primary visual cortex (inset) and projection of the first 850 brain images (green channel) acquired with serial two-photon tomography, overlaid with the UAA atlas. Scale bar = 500 μm. Right: brainrender rendering of the position of the segmented regions (red and cyan) relative to V1M (orange) and V1B (blue). Chart plot shows the proportion of the segmented regions located in V1M and V1B. **F**. Schematic of the experiment timeline. Top: Injection of an AAV carrying the fluorescent indicator GCaMP7f in the primary visual cortex (VISp) at a depth 250 μm. Middle: 2-Photon calcium imaging of layer 2/3 neurons in awake head-fixed mice, while presenting sparse noise stimuli. Example recorded trials from one neuron are shown in grey and their average response in black, aligned to the onset of stimulus. Bottom: Average Receptive Field (RF) from one neuron, shown in visual space. **G**. Left: centroids of all RFs recorded in one animal, color-coded according to their elevation in visual space. Right: centroids of all RFs recorded in one animal, color-coded according to their azimuth position in visual space. **H**. Left: brainrender rendering of the position in atlas space of all recorded cells shown in panel **G**, color-coded according to their elevation in visual space. Right: brainrender rendering of the position in atlas space of all recorded cells shown in panel **G**, color-coded according to their azimuth position in visual space.

To assess the general variability of segmentation, a centroid was computed for each segmented region and a “consensus” centroid was calculated by averaging all centroid positions (3 centroids per brain, 2 brains; n = 6). Then, each rater’s performance was assessed by calculating the Euclidean distance between their centroids and the “consensus” centroid (Figure 3B). The average distances to consensus (median = 12.3 µm, range 9.7 to 31.5 µm) were not significantly different between raters (rater 1: median = 13.4 µm, range = 10.4 to 17.4 µm; rater 2: median = 13.8 µm, range = 9.7 to 31.5 µm; rater 3: median = 10.4 µm, range = 9.9 to 17.1 µm; p>0.05, wilcoxon signed rank test). The distance to consensus didn’t exceed 31.5 µm (rater 2; Figure 3B). In addition, we assessed the intra-rater variability of region segmentation by computing the standard deviation of each rater (3 centroids per brain, 2 brains; n = 2 SDs per rater). The standard deviation never exceeded 10.5 µm (rater 2; Figure 2B).

These results show that human localisation of an injection site using the brainreg/brainreg-segment pipeline is reliable and accurate, with intra-rater accuracy typically averaging 12 µm and reliability restricted to a standard deviation of 10 µm.

We then assessed the variability in volume and shape of the segmented brain regions. The volume reported by brainreg-segment was compared between raters. The volumes of the 3 raters were all in the same order of magnitude (Figure 3C). However, volume estimation significantly differed between raters, with the exception of raters 2 and 3 (rater 1: median = 0.019 mm^3^, range = 0.017 to 0.022 mm^3^; rater 2: median = 0.021 mm^3^, range = 0.020 to 0.030 mm^3^; rater 3: median = 0.028 mm^3^, range = 0.026 to 0.037 mm^3^; p<0.05, wilcoxon signed rank test; Figure 3C). Since these raters nevertheless estimated a similar position for their respective brain segmentation centroids, the differences in volume reflects a difference in judgement of injection borders (see discussion).

The overlap of the brain segmentations was also explored by scoring each rater against a “consensus” segmented region. The consensus segmented region was taken as brain tissue (voxels) that were included in at least 50% of the raters’ segmentations. The overlap of each segmented region with the consensus was then computed in the form of a DICE score (Dice, 1945; Figure 3C). The DICE scores were not significantly different between raters (rater 1: median = 0.86, range = 0.83 to 0.90; rater 2: median = 0.92, range = 0.83 to 0.95; rater 3: median = 0.85, range = 0.80 to 0.94; p>0.05, wilcoxon signed rank test; Figure 3C). The DICE scores didn’t fall below 0.80 (rater 3), which indicates a high degree of overlap between the raters’ segmented regions and the “consensus” region (Figure 3C).

Next, the advantage of manual segmentation of injection sites using brainreg-segment was further demonstrated using two extra features. First, the brain segmentation was subdivided to reflect its spread through different brain regions. In this case, the injection in both brains were mainly contained in layers 6a and 6b (VISp6a and VISp6b), reflecting the expression pattern of the cre+ cells in the Ntsr1-cre mouse line (Figure 3D). Second, we used the UAA (Figure 1A) to characterise the injection spread according to the V1M/V1B border. We found that the injections in both brains were mainly located in V1M (83%), but also spread to V1B (Figure 3E).

Finally, we demonstrated the full potential of brainreg by functionally characterizing individual neurons in the injection site and bridging functional properties to anatomical localization. To achieve this, a glass pipette containing a solution of adenovirus carrying a construct expressing GCaMP7f was inserted in the primary visual cortex, a small volume of the solution was injected at a depth of 250 µm (layer 2/3) and a chronic cranial window was created. Sixteen days later, the GCaMP7f+ cells were imaged under a two-photon microscope and analysed using suite2p (Pachitariu et al., 2017) and their respective receptive fields (RFs) mapped (Figure 3F). The centroid of each RF was computed in visual space and color-coded according to their elevation/azimuth position (Figure 3G and H). At the end of the experiment, the animal was perfused, the brain harvested and imaged using serial two-photon tomography. The images were processed through the brainreg pipeline using the CCFv3 25 µm resolution atlas (Figure 3G). The position of each functionally imaged cell was extracted from suite2p and positioned in atlas space relative to the centroid of the GCaMP7f injection site, which was computed using the brainreg-segment “Region segmentation” feature (Figure 3H). The color coding in visual space (Figure 3G) was used to demonstrate the primary visual cortex’ retinotopic organisation of RFs (Figure 3H).

## Discussion

Traditionally, 2D histology has been used to map structures such as implanted devices within the brain, and more recently, semi-automated methods have been developed to streamline the process (Shamash et al., 2018; Peters, 2021). However these methods remain time-consuming, especially if many tissue sections must be processed (e.g. for large, or multiple implants). 3D whole-brain approaches have been developed using MRI or CT (Borg et al., 2015; Rangarajan et al.; 2016, Király et al.; 2020; Kollo et al., 2020) which permit *in-vivo* brain mapping. However, these methods are not compatible with fluorescence imaging and so can not be used to map the distribution of fluorescent proteins such as GFP.

Here we show the implementation of two integrated open-source software tools, brainreg and brainreg-segment, and we tested their implementation for localising high-density probe recording sites and fluorescence-expressing cells following the injection of recombinant adeno-associated virus carrying cre-dependent fluorescent proteins into the brain of cre reporter mice. We asked 3 neuroscientists to perform the same tasks on the same brains using the pipeline, which allowed us to establish overall and intra-rater accuracy and reliability.

Overall, we found that brainreg/brainreg-segment allowed accurate and reliable probe track tracing. Compared to “ground-truth” depth, the average accuracy error was less than 70 µm, comparable to what has been recently reported (Liu et al., 2020). In addition, we found that the reliability of raters, as measured by their standard deviations of the tracings of the same track, was less than 18 µm — but we also found that intra-rater reliability was rater-dependent. These results suggest that to reach reliable results, probe track tracing should be performed several times for each probe track and electrode localisation should be determined using an average track trace.

An interesting result of our track tracing was the systematic underestimation of probe track length by all raters (excluding one outlier brain). This reflects the difficulty of detecting accurately the DiI signal at the tip of the probe, which results in experimenters tracing shorter tracks. This issue, however, can be overcome by electrolytic lesions at the tip of the probe, thus marking the exact position of the tip (Senzai et al., 2019).

The localization of injection centroids using the brainreg/brainreg-segment pipeline was also accurate and reliable. On average, the raters centroids were 12.3 µm away from “consensus” and their reliability, assessed by the repeats in segmentation for each brain, was less than 10 µm. In addition, all raters’ segmentations had a high DICE score when compared to “consensus”, further indicating high accuracy and reliability. However, the volumes of the segmented brain regions differed according to the rater. Since the raters “agreed” with the centroid position, but “disagreed” with the injection volumes (i.e. the extent of injection spread), this result suggests that raters have defined differently the region borders. This limit can be overcome, in future brain segmentations, by agreeing to a standard fluorescent threshold for inclusion or exclusion in the segmented region.

To our knowledge, there has been a single report of this type of analysis using whole-brain microscopy. Liu et al. (2020) used LSFM to image Neuropixels probes and a semi-automated approach to align the images to an atlas (CCFv3). They manually traced probe positions, and used photostimulation of small axonal projections as a ground-truth. They compared the results of their neuroanatomical approach to ground-truth, reporting a mean accuracy of 70 µm. While this approach is comprehensive, and the accuracy is comparable to that reported here, it is specific to linear electrophysiological probes and the software is not yet publicly available.

One of the main advantages of our approach is flexibility. The software is illustrated with specific examples of electrophysiological probes and virus injections — including the segmentation, localization and functional annotation of cells that had previously been imaged using a calcium indicator. However the software is designed to segment multiple structures of any size and shape. The tool can be used to segment brain structures and fibre tracts, pathological or experimenter-made lesions, or recording/stimulation devices such as optical fibres. Fluorescence microscopy allows multiple types of structures to be labelled, imaged and segmented within a single animal. Accurate mapping of labelled neuronal populations along with recording and stimulation devices will allow for a better understanding of the complex data that modern neuroscience often relies upon.

Our software is built upon open-source Python data analysis and visualisation tools (Van Der Walt et al., 2014; Harris et al., 2020; Virtanen et al., 2020; Sofroniew et al., 2021) and continues to be developed. It is part of the BrainGlobe suite of computational neuroanatomy tools, built upon the BrainGlobe Atlas API (Claudi et al., 2020) to provide compatibility with multiple brain atlases now, and new atlases in the future. All the tools in the BrainGlobe suite are compatible with these atlases, and so the results of brainreg-segment can be directly compared with results from other software, for example cellfinder (Tyson et al., 2021), to compare the distribution of individually labelled cells to a bulk injection site.

One limitation of this software is manual segmentation. This approach was chosen as it is relatively fast — segmentation times depend on the structure being segmented, but are typically from seconds to a few (e.g. five) minutes. It also requires no optimisation, and it is relatively consistent between raters. However, as whole-brain microscopy becomes easier and cheaper, and large multi-laboratory efforts become more common in neuroscience (e.g. Abbot et al. 2017), fully automated methods could improve the reliability and reproducibility of results. In the future, we aim to provide support for automated segmentation methods (Breiman 2001; Ronneberger et al., 2015), along with additional registration algorithms (Avants et al., 2011; Klein et al., 2010) and more atlases via the BrainGlobe Atlas API.

## Data and software availability

The software outlined in this manuscript is available as part of the BrainGlobe suite of computational neuroanatomy tools. The software is open-source, written in Python 3 and runs on standard desktop computing hardware. Brainreg is fully supported on Linux and Windows, and brainreg-segment on Linux, Windows and macOS. Source code is available at github.com/brainglobe/brainreg and github.com/brainglobe/brainreg-segment. Documentation and tutorials are available at docs.brainglobe.info.

## Acknowledgements

We thank Molly Strom for assistance with virus development and sample preparation. This work was supported by grants from the Gatsby Charitable Foundation (GAT3361) and Wellcome Trust (090843/F/09/Z and 214333/Z/18/Z) to T.W.M.

## Materials and methods

### Surgical procedures

All experiments were performed on 10-18 week old C57BL/6, B6.FVB(Cg)-Tg(Ntsr1-cre)Gn220Gsat/Mmucd (Ntsr1-cre), or B6.Cg-GT(ROSA)26 Sortm14(CAG-tdTomato)Hze/J crossed to Gad2tm2(cre)Zjh/J male mice in accordance with the UK Home Office regulations (Animal Welfare Act 2006) and approved by the establishments Animal Welfare and Ethical Review Board. All surgical procedures, including the implantation of head plates, craniotomies, cranial windows and virus injections, were carried out under isoflurane (2%-5%) and after carprofen (5mg/kg, s.c.) had been administered.

### Extracellular recordings

For extracellular recordings, C57BL/6 mice (10-18 weeks old) were used. Mice were anesthetized under isoflurane (2%-5%), implanted with a head plate and allowed to recover for 48 hr. Then, animals were anesthetized under isoflurane (2%-5%) and a small craniotomy (1 × 1 mm) was drilled over the primary visual cortex. The craniotomy was then sealed with silicon (kwik-cast) and the animals were allowed to recover for 2 hr. The head was then positioned with a pitch angle of 30 degrees (nose down). The silicon probe was first coated with DiI (Molecular Probes, Thermo Fisher Scientific, USA). The probe was then positioned perpendicular to the brain surface and the tip inserted to 1750 or 2000 µm from the pial surface. The silicon probe (IMEC, Belgium) data was acquired using an FPGA card (KC705, Xilinx, USA) and SpikeGLX (https://github.com/billkarsh/SpikeGLX) and filtered at 300 Hz.

To determine the middle of layer 5 electrophysiologically, we used the depth profile of the high-frequency LFP power (Senzai et al., 2019). Briefly, for each non-reference channel on the probe, the average power in the frequency range 500-1250 Hz was computed (MATLAB, The MathWorks Inc., bandpower function). Reference channels were interpolated (MATLAB interp1), and the depth profile was smoothed across 10 channels (MATLAB smooth, which implements a moving average) to reduce variability across channels. The location, in the cortex, of the peak in this depth profile was chosen as the middle of layer 5 (mid-VISp5).

### Virus injections

For virus injections, the same procedure was followed as described in Vélez-Fort et al., 2014. Briefly, Ntsr1-cre mice (10-15 weeks old) were anaesthetized under isoflurane (2%-5%) and craniotomies performed. Cre-dependent AAV viruses encoding the fluorescent protein GFP were injected in layer 6 of the primary visual cortex. After injections, the craniotomy was sealed with silicon (kwik-cast) and animals were allowed to recover for 2-3 weeks. Viruses were delivered at a rate of 1-2 nl/s using Nanoject III (Drummond Scientific, USA).

For *in vivo* GCaMP7f imaging, mice were anaesthetized with isoflurane (2%-5%) and implanted with a headplate. A craniotomy (3 mm) was drilled over the primary visual cortex, and 100 nl of a virus encoding GCaMP7f (AAV1-syn-jGCaMP7f-WPRE, titre: 4.75×10^12^ ml^-1^) was injected in layer 2/3 of the primary visual cortex (Bregma: −2.7 mm, ML: −2.58 mm, depth: 0.25 mm) at a rate of 1 nl/s. The craniotomy was sealed by implanting a 3 mm diameter round glass coverslip (#1, Harvard Apparatus, USA) to create a chronic cranial window.

### Two-photon calcium imaging

Prior to imaging experiments, mice were handled and accustomed to head fixation for 3 days. GCaMP7f fluorescence was imaged 16 days after viral injection in awake head-fixed mice using a two-photon moveable objective microscope (Sutter Instruments, USA) coupled to a Ti-Sapphire laser (Insight Deepsee, Spectra Physics, USA) tuned to 930 nm. Neurons located ∼200 µm below the surface of the brain were visualised with a 16x 0.8 NA water-immersion objective (Nikon, Japan) and GaAsP PMTs (Hamamatsu Photonics, Japan) in a 390×390 µm^2^ field of view and images (512 × 512 pixels) were acquired with a resonant scanner (8kHz, Cambridge Technology) at a frame rate of 30 Hz. To minimize photodamage, the excitation laser power was adjusted to the lowest intensity required for the imaging depth.

### Visual stimulation

Visual stimuli for receptive field (RF) mapping were generated in MATLAB using the Psychophysics Toolbox (Brainard, 1997) and presented on an LCD monitor (Samsung C24FG73FQU, 120 Hz refresh rate) positioned 20 cm from the right eye at approximately 45° to the long axis of the animal, such that it covered ∼105° (azimuth) × 75° (elevation) of visual105° (azimuth) × 75° (elevation) of visual space. Stimuli consisted of black and white squares on a grey background, presented one at a time (1Hz) in a randomised order, at one of 120 positions (12 × 10 matrix) covering the full surface of the monitor. Each position was repeated 10 times. The backlight of the monitor was synchronized to one turn-around point of the resonant scanner (where data is not acquired), so that the monitor flickered at 8kHz and light from the screen is only on in between the scanning of two subsequent lines. A photodiode was positioned over the upper-left corner of the monitor and data was acquired in parallel with imaging to provide an accurate stimulus timestamp.

### Receptive field analysis

Preprocessing of all raw calcium movie data including image registration and automated cell detection was done in suite2p (Pachitariu et al., 2017). All subsequent analysis was performed with custom scripts written in MATLAB. Neuropil subtraction to correct for contamination of the ROI calcium traces by surrounding neuropil and calculation of ΔF/F was computed as described in de Vries et al., 2020. The response to each individual patch was taken as the mean dF/F during the 1s stimulus presentation minus the mean dF/F during the previous 1s (baseline). In the following, responses to black and white stimuli were analysed separately. To assess whether ROIs had a significant RF, responses to each patch position were grouped and a Kruskal-Wallis test was performed (p < 0.05). Only ROIs with significant RFs were subsequently analysed. To compute the RF centroid, the median response to each stimulus position was taken to create a raw 12 × 10 RF map. These RF maps were smoothed (imgaussfilt) and upsampled by a factor of 10 (imresize). Upsampled RFs were then thresholded (*threshold* = max(RF) - *std*(RF)) and the centroid of the largest subdomain after thresholding was taken as the RF centroid.

### Serial two-photon imaging

Following silicon probe recordings or virus injections, animals were deeply anaesthetized and transcardially perfused with cold phosphate buffer (PB, 0.1 M) followed by 4% paraformaldehyde (PFA) in PB (0.1 M) and brains left overnight in 4% PFA at 4 °C. Brains were then embedded in 4% agar and imaged using serial two-photon tomography (Ragan et al., 2012), using a commercially available system as previously described (Vélez-Fort et al., 2014) or a custom system controlled by ScanImage (v5.6, Vidrio Technologies, USA) using BakingTray (Campbell, 2020a). Images were acquired as tiles, and stitched using a custom FIJI (Schindelin et al., 2012) plugin (modified from Preibisch et al., 2009) or StitchIt (Campbell et al., 2020b). For Neuropixels probe tracking, images were acquired with 20μm axial sampling with 4.6 × 4.6μm pixels. For GFP-injection mapping, images were acquired with 1 × 1μm pixels and 5μm axial sampling, before downsampling to 5μm isotropic sampling. For GCaMP7f-injection mapping, images were acquired with 2.23 × 2.23 μm pixels and 10 μm axial sampling.

### Image registration

Image registration was carried out using brainreg, which is a Python port of the automatic mouse atlas propagation (aMAP) software (Niedworok et al., 2016), and is a wrapper around niftyreg (Modat et al., 2010) to carry out the registration. Images were registered to an atlas, either the Allen Mouse Brain Common Coordinate Framework version 3 (Wang et al., 2020) or the Unified Anatomical Atlas (Chon et al., 2019). All atlas data was provided by the BrainGlobe Atlas API (Claudi et al., 2020).

For registration, the sample image data was initially downsampled to the voxel spacing of the atlas used (either 10, 25, 50 or 100 μm isotropic), and reoriented to align with the atlas orientation using bg-space (Petrucco & Tyson, 2021). The downsampled image and the atlas reference images were then filtered using scikit-image (Van Der Walt et al., 2014) and SciPy (Virtanen et al., 2020), in order to remove-high frequency noise (greyscale opening and flat-field correction). Registration was performed in two steps, firstly images were aligned using an affine transform using the NiftyReg reg_aladin command (Ourselin et al., 2001). Non-linear registration was then performed with reg_f3d (Modat et al., 2010). The transformation from atlas space to sample space was then applied to the atlas annotations image (and an atlas hemispheres image) so that brain region annotations could be overlaid on the image. For analysis in atlas space, the affine transform was inverted (using reg_transform), and the non-linear transformation was performed in reverse (from sample to atlas).

### Manual segmentation and region analysis

To manually segment structures within the brain, a graphical user interface was developed using Qt (www.qt.io/ via qtpy https://github.com/spyder-ide/qtpy) and napari (Sofroniew et al., 2021). This tool (brainreg-segment) allows 1, 2, and 3-dimensional structures to be segmented, analysed (e.g. distribution within brain regions) and summarised (e.g. centroids of volumes).

A full user-guide for brainreg-segment is available at docs.brainglobe.info/brainreg-segment/introduction, but a summary of the analysis process is as follows. Upon starting the plugin, the user is presented with an option to load data previously registered with brainreg. The user can load the entire brainreg output directory, which loads the appropriate atlas data and image metadata. The analysis can be carried out in the coordinate space of the atlas, the sample, or alternatively, the atlas alone can be loaded. All analysis in this manuscript was performed in atlas space.

For segmentation of 1D structures (e.g. Neuropixels probes), there is a “Track tracing” button which loads a new panel. Selecting the “Add track” button allows the user to follow the path of the structure by adding points along its length. These points can then be fitted (“Trace tracks” button) using spline interpolation and the user can choose the order of the spline fit, a smoothing factor (how closely or not to fit the points) and the number of points along the track to analyse. In this manuscript, we used a linear fit (order 1), smoothing of 0.1 and 1750 or 2000 points, which corresponds to one sampling point per micron. The analysis of this fit results in a CSV file detailing the brain region at each of the sampling points along the length of the structure.

For segmentation of 2/3D structures (e.g. injection sites), there is a “Region segmentation” button which loads a different panel. Selecting the “Add track button” allows the user to segment the structure by “painting” onto the image in successive 2D planes. The spatial distribution of the structure can then be analysed (“Analyse regions” button), and various parameters can be saved, such as the volume of the structure in each brain region, and summary statistics such as the centroid of each volume.

There is no limit to the number of structures that can be segmented using this tool, and there is a button to save the structures (to be reloaded at a later date). If analysis is performed in atlas space, the structures can be directly exported in a brainrender (Claudi et al., 2021) compatible format (.npy or .obj) for visualisation in 3D.

### Computing hardware

For timing of registration (Figure 1G), analyses were performed on two machines, a desktop workstation and a laptop. The workstation ran Ubuntu 16.04, and had 384GB RAM and dual Intel Xeon 6132 14 core (28 thread) CPUs, but brainreg was limited to 6 (12) cores for registration. The laptop ran Ubuntu 18.04 and had 16GB RAM, and a single Intel i7-8565U (4 core, 8 thread) CPU. In both cases, data were stored on an M.2 NVMe drive.

### Data analysis and visualisation

To compare segmented injection variability, a consensus injection image was calculated from all nine (three repeats by three experts) manual segmentations. Voxels which were found in over half of the segmentations (i.e. at least five) made up the binary mask of the consensus image. Individual expert segmentations were then compared to this consensus by calculating the DICE (Dice, 1945) score.

Plots were generated using MATLAB (The MathWorks Inc.) and image visualisation was performed using napari and brainrender.

## References

Abbott, L.F., Angelaki, D.E., Carandini, M., Churchland, A.K., Dan, Y., Dayan, P., Deneve, S., Fiete, I., Ganguli, S., Harris, K.D., Häusser, M., Hofer, S., Latham, P.E., Mainen, Z.F., Mrsic-Flogel, T., Paninski, L., Pillow, J.W., Pouget, A., Svoboda, K., Witten, I.B., Zador, A.M., 2017. An International Laboratory for Systems and Computational Neuroscience. Neuron 96, 1213–1218. https://doi.org/10.1016/j.neuron.2017.12.013

Adelsberger, H., Garaschuk, O., Konnerth, A., 2005. Cortical calcium waves in resting newborn mice. Nat. Neurosci. 8, 988–990. https://doi.org/10.1038/nn1502

Arenkiel, B.R., Peca, J., Davison, I.G., Feliciano, C., Deisseroth, K., Augustine, G.J.J., Ehlers, M.D., Feng, G., 2007. In Vivo Light-Induced Activation of Neural Circuitry in Transgenic Mice Expressing Channelrhodopsin-2. Neuron 54, 205–218. https://doi.org/10.1016/j.neuron.2007.03.005

Avants, B.B., Tustison, N.J., Song, G., Cook, P.A., Klein, A., Gee, J.C., 2011. A reproducible evaluation of ANTs similarity metric performance in brain image registration. Neuroimage 54, 2033–2044. https://doi.org/10.1016/j.neuroimage.2010.09.025

Berényi, A., Somogyvári, Z., Nagy, A.J., Roux, L., Long, J.D., Fujisawa, S., Stark, E., Leonardo, A., Harris, T.D., Buzsáki, G., 2014. Large-scale, high-density (up to 512 channels) recording of local circuits in behaving animals. J. Neurophysiol. 111, 1132–1149. https://doi.org/10.1152/jn.00785.2013

Boyden, E.S., Zhang, F., Bamberg, E., Nagel, G., Deisseroth, K., 2005. Millisecond-timescale, genetically targeted optical control of neural activity. Nat. Neurosci. 8, 1263–1268. https://doi.org/10.1038/nn1525

Borg, J.S., Vu, M.A., Badea, C., Badea, A., Johnson, G.A., Dzirasa, K., 2015. Localization of metal electrodes in the intact rat brain using registration of 3D microcomputed tomography images to a magnetic resonance histology atlas. eNeuro 2, 1–14. https://doi.org/10.1523/ENEURO.0017-15.2015

Brainard, D. H., 1997. The Psychophysics Toolbox. Spatial Vision, 10(4), 433–436. http://www.ncbi.nlm.nih.gov/pubmed/9176952

Breiman, L., 2001. Random Forests. Machine Learning 45, 5–32. https://doi.org/10.1023/A:1010933404324

Campbell, R.A.A., 2020a. BakingTray: Serial-section automated anatomy extension for ScanImage. https://doi.org/doi.org/10.5281/zenodo.3631610

Campbell R.A.A., Blot, A., lguerard., 2020b. StitchIt: Stitching of large tiled datasets. http://doi.org/10.5281/zenodo.3941901

Chung, K., Wallace, J., Kim, S., Kalyanasundaram, S., Andalman, A.S., Davidson, T.J., Mirzabekov, J.J., Zalocusky, K.A., Mattis, J., Denisin, A.K., Pak, S., Bernstein, H., Ramakrishnan, C., Grosenick, L., Gradinaru, V., Deisseroth, K., 2013. Structural and molecular interrogation of intact biological systems. Nature 497, 332–337. https://doi.org/10.1038/nature12107

Chon, U., Vanselow, D.J., Cheng, K.C., Kim, Y., 2019. Enhanced and Unified Anatomical Labeling for a Common Mouse Brain Atlas. Nat. Commun. 5067. https://doi.org/10.1101/636175

Claudi, F., Petrucco, L., Tyson, A.L., Branco, T., Margrie, T.W., Portugues, R., 2020. BrainGlobe Atlas API: a common interface for neuroanatomical atlases. J. Open Source Softw. 5, 2668. https://doi.org/10.21105/joss.02668

Claudi F., Tyson A.L., Petrucco L., Margrie T.W., Portugues R., Branco T., 2021. Visualizing anatomically registered data with brainrender. eLife 10:e65751 https://doi.org/10.7554/eLife.65751

Cooper, S., Daniel, P.M., Whitteridge, D., 1953. Nerve impulses in the brainstem of the goat. Short latency responses obtained by stretching the extrinsic eye muscles and the jaw muscles. J. Physiol. 120, 471–490. https://10.1113/jphysiol.1953.sp004912

de Vries, S.E.J., Lecoq, J.A., Buice, M.A., Groblewski, P.A., Ocker, G.K., Oliver, M., Feng, D., Cain, N., Ledochowitsch, P., Millman, D., Roll, K., Garrett, M., Keenan, T., Kuan, L., Mihalas, S., Olsen, S., Thompson, C., Wakeman, W., Waters, J., … Koch, C., 2020. A large-scale standardized physiological survey reveals functional organization of the mouse visual cortex. Nat. Neurosci. 23(1), 138–151. https://doi.org/10.1038/s41593-019-0550-9

Dice, L.R., 1945. Measures of the Amount of Ecologic Association Between Species. Ecology 26, 297–302. https://doi.org/10.2307/1932409

Dodt, H., Leischner, U., Schierloh, A., 2007. Ultramicroscopy: three-dimensional visualization of neuronal networks in the whole mouse brain. Nat. Methods 4, 331–336. https://doi.org/10.1038/NMETH1036

Economo, M.N., Clack, N.G., Lavis, L.D., Gerfen, C.R., Svoboda, K., Myers, E.W., Chandrashekar, J., 2016. A platform for brain-wide imaging and reconstruction of individual neurons. eLife 5, 1–22. https://doi.org/10.7554/eLife.10566

Friedmann, D., Pun, A., Adams, E.L., Lui, J.H., Kebschull, J.M., Grutzner, S.M., Castagnola, C., Tessier-Lavigne, M., Luo, L., 2020. Mapping mesoscale axonal projections in the mouse brain using a 3D convolutional network. Proc. Natl. Acad. Sci. U. S. A. 117, 11038–11047. https://doi.org/10.1073/pnas.1918465117

Furth, D., Vaissiere, T., Tzortzi, O., Xuan, Y., Martin, A., Lazaridis, I., Spigalon, G., Fisone, Gi., Tomer, R., Deisseroth, K., Carlen, M., Miller, C.A., Rumbaugh, G., Meletis, K., 2018. An interactive framework for whole-brain maps at cellular resolution. Nat. Neurosci. 21, 139–149. https://doi.org/10.2514/6.2011-3838

Gong, H., Zeng, S., Yan, C., Lv, X., Yang, Z., Xu, T., Feng, Z., Ding, W., Qi, X., Li, A., Wu, J., Luo, Q., 2013. Continuously tracing brain-wide long-distance axonal projections in mice at a one-micron voxel resolution. Neuroimage 74, 87–98. https://doi.org/10.1016/j.neuroimage.2013.02.005

Goubran, M., Leuze, C., Hsueh, B., Aswendt, M., Ye, L., Tian, Q., Cheng, M.Y., Crow, A., Steinberg, G.K., McNab, J.A., Deisseroth, K., Zeineh, M., 2019. Multimodal image registration and connectivity analysis for integration of connectomic data from microscopy to MRI. Nat. Commun. 10, 1–17. https://doi.org/10.1038/s41467-019-13374-0

Grienberger, C., Konnerth, A., 2012. Imaging Calcium in Neurons. Neuron 73, 862–885. https://doi.org/10.1016/j.neuron.2012.02.011

Harris, C.R., Millman, K.J., van der Walt, S.J., Gommers, R., Virtanen, P., Cournapeau, D., Wieser, E., Taylor, J., Berg, S., Smith, N.J., Kern, R., Picus, M., Hoyer, S., van Kerkwijk, M.H., Brett, M., Haldane, A., del Río, J.F., Wiebe, M., Peterson, P., Gérard-Marchant, P., Sheppard, K., Reddy, T., Weckesser, W., Abbasi, H., Gohlke, C., Oliphant, T.E., 2020. Array programming with NumPy. Nature 585, 357–362. https://doi.org/10.1038/s41586-020-2649-2

Hoops, D., Weng, H., Shahid, A., Skorzewski, P., Janke, A.L., Lerch, J.P., Sled, J.G., 2021. A fully segmented 3D anatomical atlas of a lizard brain. Brain Struct. Funct. https://doi.org/10.1007/s00429-021-02282-z

Iqbal, A., Sheikh, A., Karayannis, T., 2019. DeNeRD: high-throughput detection of neurons for brain-wide analysis with deep learning. Sci. Rep. 9, 1–13. https://doi.org/10.1038/s41598-019-50137-9

Jin, M., Nguyen, J.D., Weber, S.J., Mejias-Aponte, C.A., Madangopal, R., Golden, S.A., 2019. SMART: An open source extension of WholeBrain for iDISCO+ LSFM intact mouse brain registration and segmentation. bioRxiv. https://doi.org/10.1101/727529

Jun, J.J., Steinmetz, N.A., Siegle, J.H., Denman, D.J., Bauza, M., Barbarits, B., Lee, A.K., Anastassiou, C.A., Andrei, A., Aydin, Ç., Barbic, M., Blanche, T.J., Bonin, V., Couto, J., Dutta, B., Gratiy, S.L., Gutnisky, D.A., Häusser, M., Karsh, B., … Harris, T.D., 2017. Fully integrated silicon probes for high-density recording of neural activity. Nature 551, 232–236. https://doi.org/10.1038/nature24636

Kenney, J.W., Steadman, P.E., Young, O., Shi, M.T., Polanco, M., Dubaishi, S., Mueller, T., Frankland, P.W., 2021. AZBA: A 3D Adult Zebrafish Brain Atlas for the Digital Age. bioRxiv. https://doi.org/10.1101/2021.05.04.442625

Király, B., Balázsfi, D., Horváth, I., Solari, N., Sviatkó, K., Lengyel, K., Birtalan, E., Babos, M., Bagaméry, G., Máthé, D., Szigeti, K., Hangya, B., 2020. In vivo localization of chronically implanted electrodes and optic fibers in mice. Nat. Commun. 11. https://doi.org/10.1038/s41467-020-18472-y

Kirst, C., Skriabine, S., Vieites-Prado, A., Topilko, T., Bertin, P., Gerschenfeld, G., Verny, F., Topilko, P., Michalski, N., Tessier-Lavigne, M., Renier, N., 2020. Mapping the Fine-Scale Organization and Plasticity of the Brain Vasculature. Cell 1–16. https://doi.org/10.1016/j.cell.2020.01.028

Klein, S., Staring, M., Murphy, K., Viergever, M.A., Pluim, J.P.W., 2010. Elastix: A toolbox for intensity-based medical image registration. IEEE Trans. Med. Imaging 29, 196–205. https://doi.org/10.1109/TMI.2009.2035616

Kollo, M., Racz, R., Hanna, M.E., Obaid, A., Angle, M.R., Wray, W., Kong, Y., Müller, J., Hierlemann, A., Melosh, N.A., Schaefer, A.T., 2020. CHIME: CMOS-Hosted in vivo Microelectrodes for Massively Scalable Neuronal Recordings. Front. Neurosci. 14, 1–13. https://doi.org/10.3389/fnins.2020.00834

Liu, L.D., Chen, S., Economo, M.N., Li, N., Svoboda, K., 2020. Accurate localization of linear probe electrodes across multiple brains. bioRxiv. https://doi.org/10.1101/2020.02.25.965210

Mano, T., Murata, K., Kon, K., Shimizu, C., Ono, H., Yamada, R.G., Miyamichi, K., Susaki, E.A., Ueda, H.R., 2020. CUBIC-Cloud : An Integrative Computational Framework Towards Community-driven Whole-Mouse-Brain Mapping. bioRxiv. https://doi.org/10.1101/2020.08.28.271031

Mei, Y., Zhang, F., 2012. Molecular tools and approaches for optogenetics. Biol. Psychiatry 71, 1033–1038. https://doi.org/10.1016/j.biopsych.2012.02.019

Modat, M., Ridgway, G.R., Taylor, Z.A., Lehmann, M., Barnes, J., Hawkes, D.J., Fox, N.C., Ourselin, S., 2010. Fast free-form deformation using graphics processing units. Comput. Methods Programs Biomed. 98, 278–284. https://doi.org/10.1016/j.cmpb.2009.09.002

Muller, E., Bednar, J.A., Diesmann, M., Gewaltig, M.O., Hines, M., Davison, A.P., 2015. Python in neuroscience. Front. Neuroinform. 9, 14–17. https://doi.org/10.3389/fninf.2015.00011

Ni, H., Feng, Z., Guan, Y., Jia, X., Chen, W., Jiang, T., Zhong, Q., Yuan, J., Ren, M., Li, X., Gong, H., Luo, Q., Li, A., 2020. DeepMapi: a Fully Automatic Registration Method for Mesoscopic Optical Brain Images Using Convolutional Neural Networks. Neuroinformatics https://doi.org/10.1007/s12021-020-09483-7

Niedworok, C.J., Brown, A.P.Y., Jorge Cardoso, M., Osten, P., Ourselin, S., Modat, M., Margrie, T.W., 2016. AMAP is a validated pipeline for registration and segmentation of high-resolution mouse brain data. Nat. Commun. 7, 1–9. https://doi.org/10.1038/ncomms11879

Ortiz, C., Navarro, J.F., Jurek, A., Märtin, A., Lundeberg, J., Meletis, K., 2020. Molecular atlas of the adult mouse brain. Sci. Adv. 6, 1–14. https://doi.org/10.1126/sciadv.abb3446

Osten, P., Margrie, T.W., 2013. Mapping brain circuitry with a light microscope. Nat. Methods 10(6), 515–523. https://doi.org/10.1038/nmeth.2477

Ourselin, S., Roche, A., Subsol, G., Pennec, X., Ayache, N., 2001. Reconstructing a 3D structure from serial histological sections. Image Vis. Comput. 19, 25–31. https://doi.org/10.1016/S0262-8856(00)00052-4

Pachitariu, M., Stringer, C., Dipoppa, M., Schröder, S., Rossi, L. F., Dalgleish, H., Carandini, M., Harris, K. D., 2017. Suite2p: beyond 10,000 neurons with standard two-photon microscopy. BioRxiv. https://doi.org/10.1101/061507

Perens, J., Gravesen, S.C., Skytte, J.L., Roostalu, U., Dahl, A.B., Dyrby, T.B., Wichern, F., Barkholt, P., Vrang, N., Jelsing, J., Hecksher-Sørensen, J., 2020. An Optimized Mouse Brain Atlas for Automated Mapping and Quantification of Neuronal Activity Using iDISCO+ and Light Sheet Fluorescence Microscopy. Neuroinformatics https://doi.org/10.1007/s12021-020-09490-8

Peters, A.J., 2021. AP_histology, GitHub repository: https://github.com/petersaj/AP_histology

Petrucco, L., Tyson, A.L., 2021. bg-space http://doi.org/10.5281/zenodo.4552537

Preibisch, S., Saalfeld, S., Tomancak, P., 2009. Globally optimal stitching of tiled 3D microscopic image acquisitions. Bioinformatics 25, 1463–1465. https://doi.org/10.1093/bioinformatics/btp184

Ragan, T., Kadiri, L.R., Venkataraju, K.U., Bahlmann, K., Sutin, J., Taranda, J., Arganda-Carreras, I., Kim, Y., Seung, H.S., Osten, P., 2012. Serial two-photon tomography for automated ex vivo mouse brain imaging. Nat. Methods 9, 255–258. https://doi.org/10.1038/nmeth.1854

Rangarajan, J.R., Vande Velde, G., Van Gent, F., De Vloo, P., Dresselaers, T., Depypere, M., Van Kuyck, K., Nuttin, B., Himmelreich, U., Maes, F., 2016. Image-based in vivo assessment of targeting accuracy of stereotactic brain surgery in experimental rodent models. Sci. Rep. 6, 1–17. https://doi.org/10.1038/srep38058

Renier, N., Adams, E.L., Kirst, C., Wu, Z., Azevedo, R., Kohl, J., Autry, A.E., Kadiri, L., Umadevi Venkataraju, K., Zhou, Y., Wang, V.X., Tang, C.Y., Olsen, O., Dulac, C., Osten, P., Tessier-Lavigne, M., 2016. Mapping of Brain Activity by Automated Volume Analysis of Immediate Early Genes. Cell 165, 1789–1802. https://doi.org/10.1016/j.cell.2016.05.007

Renier, N., Wu, Z., Simon, D.J., Yang, J., Ariel, P., Tessier-Lavigne, M., 2014. iDISCO: A Simple, Rapid Method to Immunolabel Large Tissue Samples for Volume Imaging. Cell 159, 896–910. https://doi.org/10.1016/j.cell.2014.10.010

Renshaw, B., Forbes, A., Morison, B.R., 1940. Activity of Isocortex and Electrical Hippocampus: Electrical Studies with Micro-Electrodes. J. Neurophysiol. 3, 74–105. https://doi.org/10.1152/jn.1940.3.1.74

Ronneberger O., Fischer P., Brox T. 2015. U-Net: Convolutional Networks for Biomedical Image Segmentation. In: Navab N., Hornegger J., Wells W., Frangi A. (eds) Medical Image Computing and Computer-Assisted Intervention – MICCAI 2015. MICCAI 2015. Lecture Notes in Computer Science, vol 9351. Springer, Cham. https://doi.org/10.1007/978-3-319-24574-4_28

Schindelin, J., Arganda-Carreras, I., Frise, E., Kaynig, V., Longair, M., Pietzsch, T., Preibisch, S., Rueden, C., Saalfeld, S., Schmid, B., Tinevez, J.-Y., White, D.J., Hartenstein, V., Eliceiri, K., Tomancak, P., Cardona, A., 2012. Fiji: an open-source platform for biological-image analysis. Nat. Methods 9, 676–682. https://doi.org/10.1038/nmeth.2019

Senzai, Y., Fernandez-Ruiz, A., Buzsáki, G., 2019. Layer-Specific Physiological Features and Interlaminar Interactions in the Primary Visual Cortex of the Mouse. Neuron 101, 500-513.e5. https://doi.org/10.1016/j.neuron.2018.12.009

Seiriki, K., Kasai, A., Hashimoto, T., Schulze, W., Niu, M., Yamaguchi, S., Nakazawa, T., Inoue, K. ichi, Uezono, S., Takada, M., Naka, Y., Igarashi, H., Tanuma, M., Waschek, J.A., Ago, Y., Tanaka, K.F., Hayata-Takano, A., Nagayasu, K., … Hashimoto, H., 2017. High-Speed and Scalable Whole-Brain Imaging in Rodents and Primates. Neuron 94, 1085-1100.e6. https://doi.org/10.1016/j.neuron.2017.05.017

Shamash, P., Carandini, M., Harris, K., Steinmetz, N., 2018. A tool for analyzing electrode tracks from slice histology. bioRxiv. https://doi.org/10.1101/447995

Sofroniew, N., Lambert, T., Evans, K., Nunez-Iglesias, J., Winston, P., Bokota, G., Yamauchi, K., Solak, A.C., ziyangczi, Peña-Castellanos G., Bussonnier, M., Buckley, G., Pop, D.D., Pam, alisterburt, Hilsenstein, V., Tung, T., Hector, Freeman J., … McGovern, A., 2021. napari/napari: 0.4.8. https://doi.org/10.5281/zenodo.4747712

Song, J.H., Choi, W., Song, Y.H., Kim, J.H., Jeong, D., Lee, S.H., Paik, S.B., 2020. Precise Mapping of Single Neurons by Calibrated 3D Reconstruction of Brain Slices Reveals Topographic Projection in Mouse Visual Cortex. Cell Rep. 31, 107682. https://doi.org/10.1016/j.celrep.2020.107682

Starr, A., Wise, K.D., Csongradi, J., 1973. An Evaluation of Photoengraved Microelectrodes for Extracellular Single-Unit Recording. IEEE Trans. Biomed. Eng. BME-20, 291–293. https://doi.org/10.1109/TBME.1973.324194

Steinmetz, N.A., Koch, C., Harris, K.D., Carandini, M., 2018. Challenges and opportunities for large-scale electrophysiology with Neuropixels probes. Curr. Opin. Neurobiol. 50, 92–100. https://doi.org/10.1016/j.conb.2018.01.009

Steinmetz N.A., Aydin C., Lebedeva A., Okun M., Pachitariu M., Bauza M., Beau M., Bhagat J., Böhm C., Broux M., Chen S., Colonell J., Gardner R.J., Karsh B., Kloosterman F., Kostadinov D., Mora-Lopez C., O’Callaghan J., Park J., … Harris T.D., 2021. Neuropixels 2.0: A miniaturized high-density probe for stable, long-term brain recordings. Science. 16, 372(6539). https://doi.org/10.1126/science.abf4588

Susaki, E.A., Tainaka, K., Perrin, D., Kishino, F., Tawara, T., Watanabe, T.M., Yokoyama, C., Onoe, H., Eguchi, M., Yamaguchi, S., Abe, T., Kiyonari, H., Shimizu, Y., Miyawaki, A., Yokota, H., Ueda, H.R., 2014. Whole-brain imaging with single-cell resolution using chemical cocktails and computational analysis. Cell 157, 726–39. https://doi.org/10.1016/j.cell.2014.03.042

Todorov, M.I., Paetzold, J.C., Schoppe, O., Tetteh, G., Shit, S., Efremov, V., Todorov-Völgyi, K., Düring, M., Dichgans, M., Piraud, M., Menze, B., Ertürk, A., 2020. Machine learning analysis of whole mouse brain vasculature. Nat. Methods 17, 442–449. https://doi.org/10.1038/s41592-020-0792-1

Tomer, R., Ye, L., Hsueh, B., Deisseroth, K., 2014. Advanced CLARITY for rapid and high-resolution imaging of intact tissues. Nat. Protoc. 9, 1682–1697. https://doi.org/10.1038/nprot.2014.123

Tyson, A.L., Rousseau, C.V., Niedworok, C.J., Keshavarzi, S., Tsitoura, C., Cossell, L., Strom, M., Margrie, T.W., 2021. A deep learning algorithm for 3D cell detection in whole mouse brain image datasets. PLoS Comp. Bio. In Press

Tyson, A.L., Margrie, T.W., 2021. Mesoscale microscopy for micromammals: image analysis tools for understanding the rodent brain. arXiv. ArXiv:2102.11812

Van Der Walt, S., Schönberger, J.L., Nunez-Iglesias, J., Boulogne, F., Warner, J.D., Yager, N., Gouillart, E., Yu, T., 2014. Scikit-image: Image processing in python. PeerJ 2014, 1–18. https://doi.org/10.7717/peerj.453

Vélez-Fort, M., Rousseau, C. V., Niedworok, C.J., Wickersham, I.R., Rancz, E.A., Brown, A.P.Y., Strom, M., Margrie, T.W., 2014. The stimulus selectivity and connectivity of layer six principal cells reveals cortical microcircuits underlying visual processing. Neuron 83, 1431–1443. https://doi.org/10.1016/j.neuron.2014.08.001

Virtanen, P., Gommers, R., Oliphant, T.E., Haberland, M., Reddy, T., Cournapeau, D., Burovski, E., Peterson, P., Weckesser, W., Bright, J., van der Walt, S.J., Brett, M., Wilson, J., Millman, K.J., Mayorov, N., Nelson, A.R.J., Jones, E., Kern, R., Larson, E., … Vázquez-Baeza, Y., 2020. SciPy 1.0: fundamental algorithms for scientific computing in Python. Nat. Methods 17, 261–272. https://doi.org/10.1038/s41592-019-0686-2

Voigt, F.F., Kirschenbaum, D., Platonova, E., Pagès, S., Campbell, R.A.A., Kastli, R., Schaettin, M., Egolf, L., van der Bourg, A., Bethge, P., Haenraets, K., Frézel, N., Topilko, T., Perin, P., Hillier, D., Hildebrand, S., Schueth, A., Roebroeck, A., Roska, B., Stoeckli, E.T., Pizzala, R., Renier, N., Zeilhofer, H.U., Karayannis, T., Ziegler, U., Batti, L., Holtmaat, A., Lüscher, C., Aguzzi, A., Helmchen, F., 2019. The mesoSPIM initiative: open-source light-sheet microscopes for imaging cleared tissue. Nat. Methods 16, 1105–1106. https://doi.org/10.1038/s41592-019-0554-0

Wang, Q., Ding, S.L., Li, Y., Royall, J., Feng, D., Lesnar, P., Graddis, N., Naeemi, M., Facer, B., Ho, A., Dolbeare, T., Blanchard, B., Dee, N., Wakeman, W., Hirokawa, K.E., Szafer, A., Sunkin, S.M., Oh, S.W., Bernard, A., Phillips, J.W., Hawrylycz, M., Koch, C., Zeng, H., Harris, J.A., Ng, L., 2020. The Allen Mouse Brain Common Coordinate Framework: A 3D Reference Atlas. Cell 181, 936-953.e20. https://doi.org/10.1016/j.cell.2020.04.007

Young, D.M., Duhn, C., Gilson, M., Nojima, M., Yuruk, D., Kumar, A., Yu, W., Sanders, S.J., 2020. Whole-Brain Image Analysis and Anatomical Atlas 3D Generation Using MagellanMapper. Curr. Protoc. Neurosci. 94, e104. https://doi.org/10.1002/cpns.10

Young, D.M., Darbandi, S.F., Schwartz, G., Bonzell, Z., Yuruk, D., Nojima, M., Gole, L., Rubenstein, J., Yu, W., Sanders, S.J., 2021. Constructing and optimizing 3D atlases from 2D data with application to the developing mouse brain. Elife 10e61408. https://doi.org/10.1101/2020.04.01.017665

